# The producer cell type of HSV-1 alters the proteomic contents and infectious capacity of virions

**DOI:** 10.1101/2025.04.04.647173

**Authors:** Gary Dunn, Luke F. Domanico, Matthew P. Taylor

## Abstract

The cell that a virus replicates in i.e., the producer cell, can alter the macromolecular composition and infectious capacity of the virions that are produced. Herpes Simplex virus type 1 (HSV-1) primarily infects keratinocytes of the epidermis or oral mucosa prior to establishing latency in neurons of the peripheral nervous system, where the virus can persist for the lifetime of the host. Many cell lines that are used to amplify HSV-1 are derived from species and tissue types that are less physiologically relevant to HSV-1 disease. To understand if the producer cell type influences HSV-1 infection, we tested the infectivity of HSV-1 derived from immortalized African green monkey kidney cells (vero), immortalized human keratinocytes (HaCaT), and primary human foreskin fibroblasts (HFF-1). We observed that the producer cell type alters the capacity of HSV-1 to produce viral proteins and infectious virions from infected cells and susceptibility to inhibition of replication by interferon treatment. HaCaT-derived HSV-1 consistently exhibited enhanced replication over HFF-1 or vero-derived virus. To determine if the producer cell type changes the protein composition of virions, we performed an untargeted LC/MS-MS analysis of virions purified from each cell line. Comparison of virion associated proteins revealed quantitative differences in composition of both cellular and viral proteins including ICP0, pUL24 and pUL42. These results highlight the influence that the producer cell-type has on HSV-1 infection outcomes and suggest that cell type specific factors can alter HSV-1 and impact viral replication.

**Importance:** Approximately 67% of the human population harbors HSV-1 infection. To study HSV-1 infection, laboratories utilize several different cell lines to propagate HSV-1 for downstream experiments. The type of cell used to produce a virus, i.e. the producer cell type, can alter the macromolecular composition, immunogenicity, and infectivity of the virions that are produced across several virus families. We found that the producer cell type of HSV-1 alters virion infectivity and virion protein composition. Therefore, the producer cell type may have implications in the spread of HSV-1 and subsequent disease outcomes in humans. Our results also raise concerns about how the use of different ceil types to propagate HSV-1 may alter the outcome, interpretation, and reproducibility of experimental results.

## Introduction

Herpes Simples Virus type 1 (HSV-1) inherently relies upon the resources of a host cell to replicate and spread. HSV-1 possesses a broad tropism both *in vitro* and *in vivo*, meaning it can successfully replicate and produce infectious virions from multiple cell types. Different cell types express distinct transcriptomes and proteomes depending on multiple factors that include the function of the cell, the extracellular environment, and cell-cell signaling. Because HSV-1 infects multiple cell types, the virus must contend with these dissimilar host environments to successfully replicate. The state of the cellular environment can change fundamental aspects of viral infection and influence the macromolecular composition and infectious properties of the virions that are produced^1–27^. An example of this phenomenon has been described for the gammaherpesvirus Epstein-Barr virus (EBV)^5^. The incorporation of viral glycoprotein complexes into the viral envelope of EBV changes based on whether the virus was produced from B cells or epithelial cells, likely due to the expression of MHC II in B cells. The change in glycoprotein incorporation results in altered cellular tropism of EBV. Beyond members of the *Herpesviridae*, producer cell-dependent effects have been described for members of at least 5 other viral families, eliciting changes in cellular tropism, immune responses, and replicative capacity^27^. It is likely that producer cell-dependent effects remain to be described for a wider range of viruses.

There are many cell lines used to propagate HSV-1 for laboratory experiments that study infection and replication. Commonly used cell lines include vero, BHK-21, and HeLa among others^28^. In this paper, we will refer to the type of cell used to propagate HSV-1 as the producer cell type. The producer cell type of HSV-1 has previously been described to alter virion protein composition. Evaluation of HSV-1 virions produced from vero and HEp-2 cells contained ICP0 and ICP4, but these proteins were not detected in BHK21-derived virions^11^. Although ICP0 and ICP4 expression are critical to the establishment of successful HSV-1 infection, it is currently unknown whether their absence in virions alters virion infectivity. Indeed, the effects that the producer cell type has on HSV-1 infectivity remains understudied. To test our hypothesis that the producer cell type of HSV-1 alters virion infectivity, we propagated HSV-1 from immortalized human keratinocytes (HaCaT), African green monkey kidney cells (vero), and human foreskin fibroblasts (HFF-1). We then compared the capacity of HSV-1 from each cell line to replicate in non-neuronal cells. Our results show that the producer cell type impacts the subsequent expression of viral proteins, production of infectious virions, and the capacity to replicate in cells stimulated by interferon in all infected cell types that were tested. To begin characterizing the molecular features that lead to these producer cell-dependent differences, we performed an untargeted analysis of the proteomic composition of HSV-1 virions. We observed changes in the composition of both viral and cellular proteins present in purified virions that was producer cell type-dependent.

## Results

### Producer cell-dependent effects on genome:PFU of HSV-1

Replication often produces non-infectious virions. Despite their lack of infectivity, non-infectious virions can still influence the replication of infectious virions. Whether non-infectious virions enhance or hinder viral replication is dependent on several factors including the type of virus and the viral strain^29–32^. We hypothesized that the producer cell type of HSV-1 alters the ratio between the number of genomes and the number of plaque forming units (PFU) (genome:PFU). To test this hypothesis, we propagated HSV-1 McKrae viral stocks from HaCaT (HaCaT-HSV), vero (Vero-HSV), and HFF-1 (HFF-HSV) cells. Each viral stock was propagated in triplicate. We used a plaque assay to determine the viral titer of HaCaT-HSV, Vero-HSV, and HFF-HSV and quantitative PCR (qPCR) to determine the absolute genome content of HSV-1 derived from each producer cell type. The genome:PFU of each producer cell type was quantified specifically for the viral stocks that were used to infect target cells for the experiments described in this paper. HaCaT-HSV had the lowest genome:PFU ratio in comparison to Vero-HSV and HFF-HSV (Table 1). This data supports the interpretation that the producer cell type alters the genome:PFU of viral stocks. The subsequent experiments discussed in this paper rely upon infection of target cells using an equivalent multiplicity of infection (MOI) based on the PFU applied per cell. It is important to note that normalizing MOI to the genome content of the viral stocks rather than the PFU/cell would result in a lower PFU/cell applied to target cells that were infected with Vero-HSV or HFF-HSV relative to cells infected with HaCaT-HSV. Previous work has shown that higher MOI infections result in enhanced replication of HSV-1^33^.

**Table 1.** Producer cell-dependent effects on the HSV-1 genome to PFU ratio. The genome to PFU ratio of HSV-1 derived from HaCaT, vero, or HFF-1 cells. Genome and PFU content were determined using qPCR to quantify genome content and plaque assay to quantify PFUs, respectively, with values being represented with ± the SEM within triplicate samples. Analysis of the statistical significance was performed relative to HaCaT-HSV using unpaired t-test. ** indicates P<0.01.

| **HSV-1 producer cell** | **Average genome:PFU ratio** | **HSV-1 infectious MOI when normalized to HaCaT-HSV genome content** |
| --- | --- | --- |
| HaCaT-HSV | 4.6 ± 0.8 | 10.0 |
| Vero-HSV | 14.4 ± 1.8 ** | 3.0 |
| HFF-HSV | 17.0 ± 3.4 ** | 3.0 |

### Producer cell-dependent effects on HSV-1 protein expression

To determine whether the producer cell type alters the capacity of virions to express viral proteins, HaCaT, vero, and HFF-1 cells were infected with HSV-1 propagated from the aforementioned cell types at an MOI of 1 and 10. Infected cells were fixed at 3 hours post infection (hpi) for MOI 10 or 6 hpi for MOI 1 and 10. Detection of viral proteins was achieved by staining with a polyclonal HSV-1 antiserum (Fig. 1a). Except for 6 hpi in HaCaTs, HaCaT-HSV consistently produced more detectable viral proteins in cells when infected at an MOI of 10 (Fig. 1b). At an MOI of 1, HaCaT-HSV produced a greater quantity of viral proteins in HaCaTs but produced equivalent amounts of protein compared to Vero-HSV in vero and HFF-1 cells (Fig. 1c). HFF-HSV consistently produced the lowest quantity of detectable viral proteins in all treatment conditions and did not produce detectable amounts of protein in HFF-1 cells at MOI 1 at either time point. In the cell types that were tested, these results show that HSV-1 produced from HaCaT cells express greater amounts of detectable viral proteins compared to HSV-1 derived from vero or HFF-1 cells.

**Figure 1.**
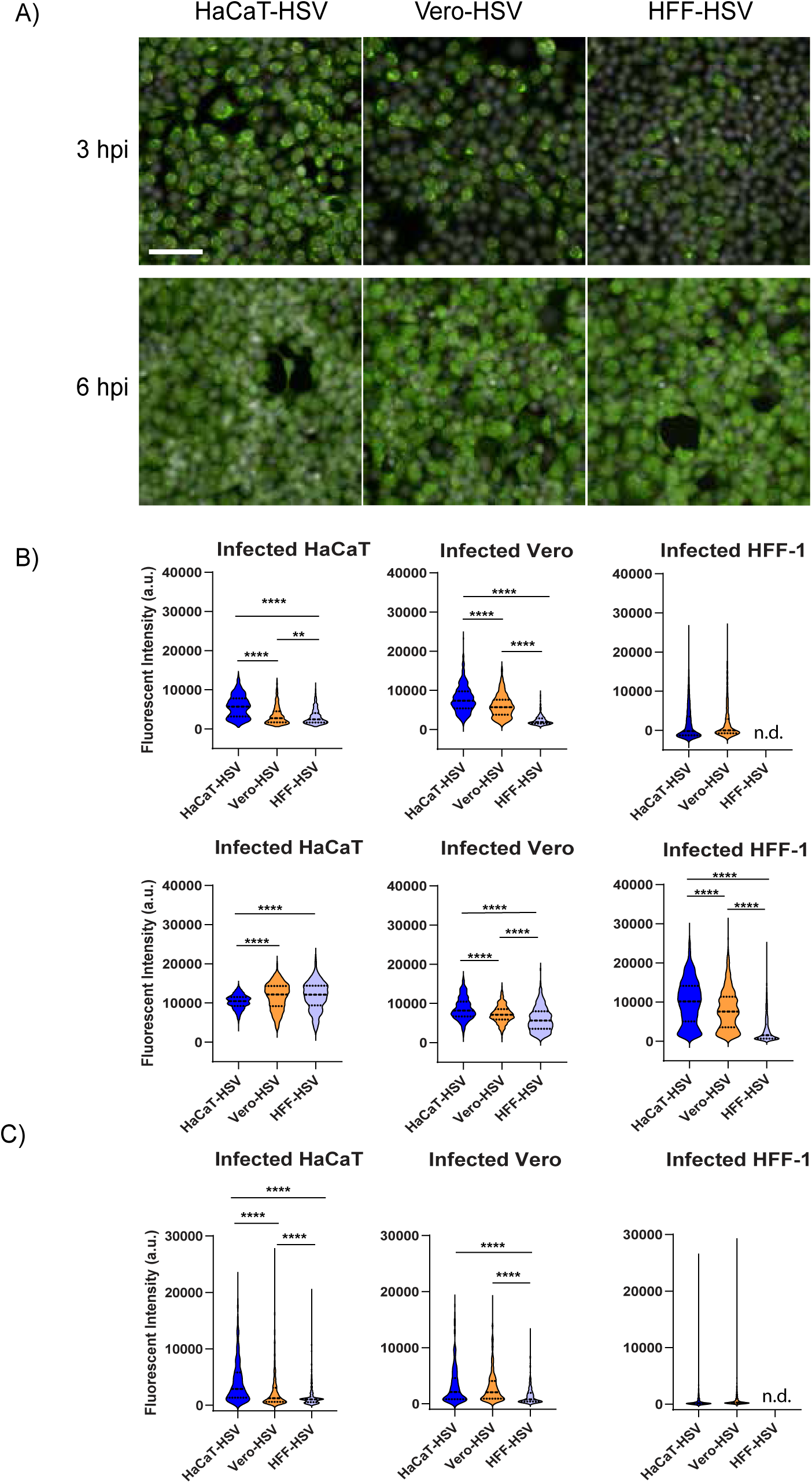
HaCaT-HSV produces more viral proteins early in infection. (A) Images of HaCaT cells infected with HSV-1 McKrae derived from HaCaT, vero, or HFF-1 cells at an MOI of 10. Infected HaCaTs were fixed at 3 and 6 hpi and stained with rabbit polyclonal HSV-1 anti-serum (green) and Hoescht (white). Equivalent exposure and pixel intensity scales are applied for each channel across all presented images. Scale bar = 200 μm. (B) Quantification of fluorescent intensity of viral proteins in HaCaT, vero, or HFF-1 cells infected with HSV-1 McKrae derived from HaCat, vero, or HFF-1 at an MOI of 10 or (C) MOI 1. Infected cells were fixed and stained as described in A at 3 hpi (top row-B and C) and 6 hpi (bottom row-B). Quantification of fluorescent intensity was performed using Image J. n.d. = no detection. Statistical comparison was performed using one-way ANOVA with Tukey’s multiple comparisons test. ** indicates p < 0.01, **** indicates p < 0.0001.

### Producer cell-dependent effects on the production of infectious HSV-1

To determine whether the producer cell type of HSV-1 alters the quantity of infectious virions produced in subsequently infected cells, HSV-1 propagated from the aforementioned cell types was inoculated onto HaCaT, vero, and HFF-1 cells at an MOI of 10. The amount of infectious virus that was produced at the indicated time points from infected cells was determined by plaque assay (Fig. 2). HaCaT-HSV consistently produced higher quantities of infectious HSV-1 in HaCaT, vero, and HFF-1 cells at 8 and 12 hpi in comparison to Vero-HSV and HFF-HSV. The output of infectious virions was roughly equivalent in all tested conditions by 24 hpi. This data shows that HaCaT-HSV has a propensity for producing higher quantities of infectious virions in infected cells.

**Figure 2.**
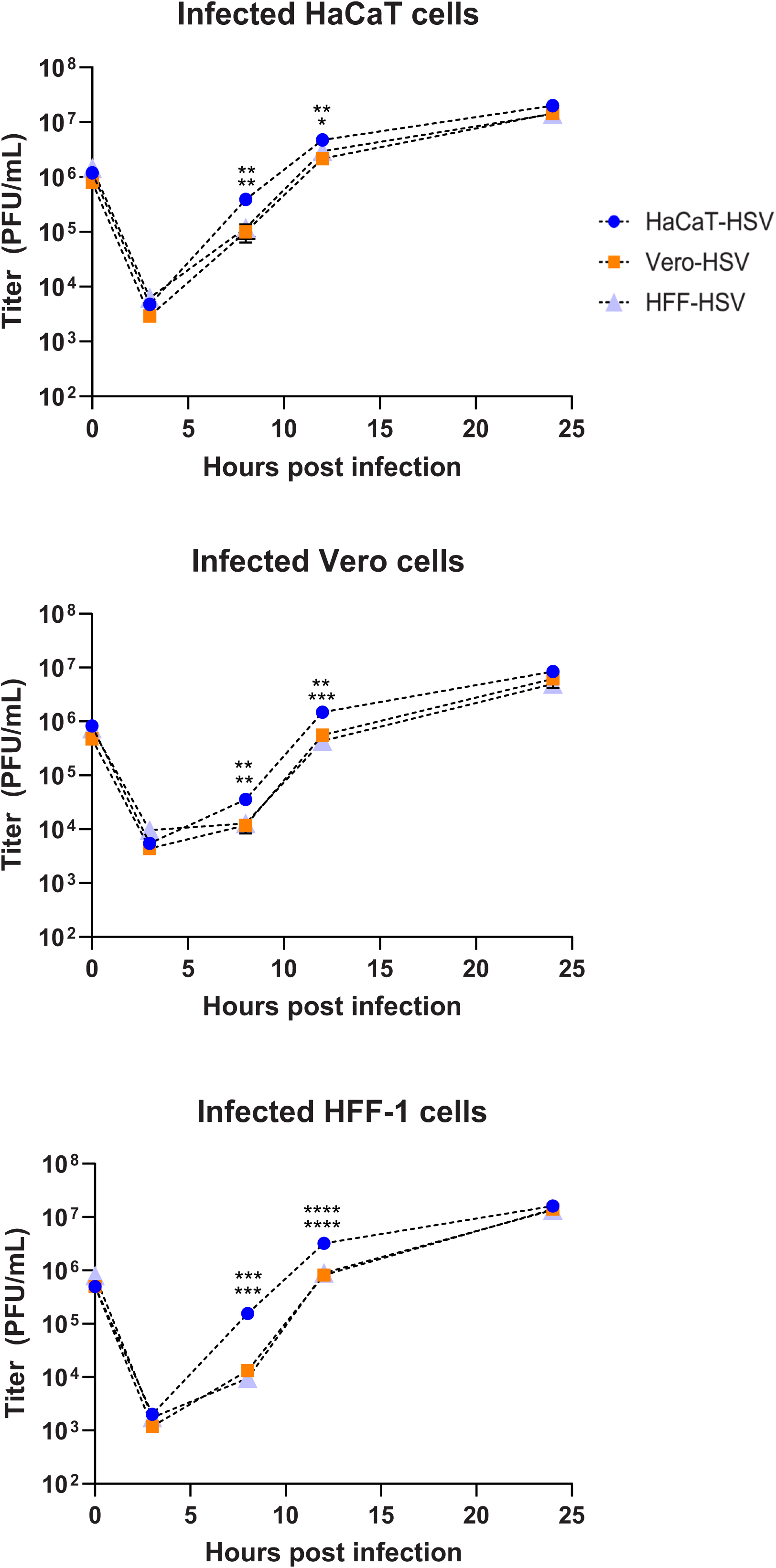
HaCaT-HSV produces higher viral titers early in infection. HaCaT, vero, and HFF-1 cells were infected with HSV-1 McKrae derived from HaCaT, vero, or HFF-1 cells at an MOI of 10. Virally infected cells and supernatant were harvested at corresponding time points after infection and titered using plaque assay on vero cells. The values represent means from three biological replicates per treatment. Error bars indicate the SEM. * indicates p < 0.05, ** indicates p < 0.01, *** indicates p < 0.001, **** indicates p<0.0001 using one-way ANOVA with Tukey’s multiple comparisons test. Statistical comparison shows HaCaT-HSV relative to Vero-HSV (top) and HaCaT-HSV relative to HFF-HSV (bottom).

### Producer cell-dependent effects on the sensitivity of HSV-1 to Type I and Type II interferon

Cells express pathogen recognition receptors (PRRs) that are capable of recognizing pathogen associated molecular patterns (PAMPs), including those associated with HSV-1 virions or replication products such as double-stranded (ds) DNA, dsRNA, or glycoproteins^34–38^. PRR recognition of viral PAMPs elicits secretion of interferons (IFN)^39^. HSV-1 expresses several viral proteins that inhibit recognition of PAMPs and downstream signaling post-PAMP recognition, but HSV-1 PAMPs eventually trigger IFN secretion from infected cells^40–43^. Over the course of viral spread, HSV-1 inevitably encounters cells that have entered an antiviral state through uptake of IFNs, including IFN-γ and IFN-β^44–47^. IFN secretion limits viral replication and spread within hosts, and HSV-1 replication is severely dampened in cells pre-treated with type I and type II IFN^45,48–60^. To test if the producer cell type influences the sensitivity of HSV-1 to IFN responses, we evaluated HSV-1 replication following pre-treatment of cells with IFN-β or IFN-γ. Target cells were pre-treated for 16 hours with IFN-β or IFN-γ (Fig. 3) prior to infection with HSV-1 McKrae propagated from the previously identified cell types. In both target cell types, IFN-β had a greater inhibitory effect on viral replication based on output viral titer in comparison to IFN-γ. Interestingly, HaCaT-HSV maintained the ability to produce higher quantities of infectious virions in comparison to Vero-HSV and HFF-HSV when infecting HaCaT and HFF-1 cells pre-treated with each interferon. Vero-HSV and HFF-HSV did not produce a statistically significant difference in the quantity of infectious virions from IFN-treated cells relative to one another. These results show that HSV-1 produced from HaCaT cells is less sensitive to the antiviral effects of type I and II IFN in the cell types that were infected.

**Figure 3.**
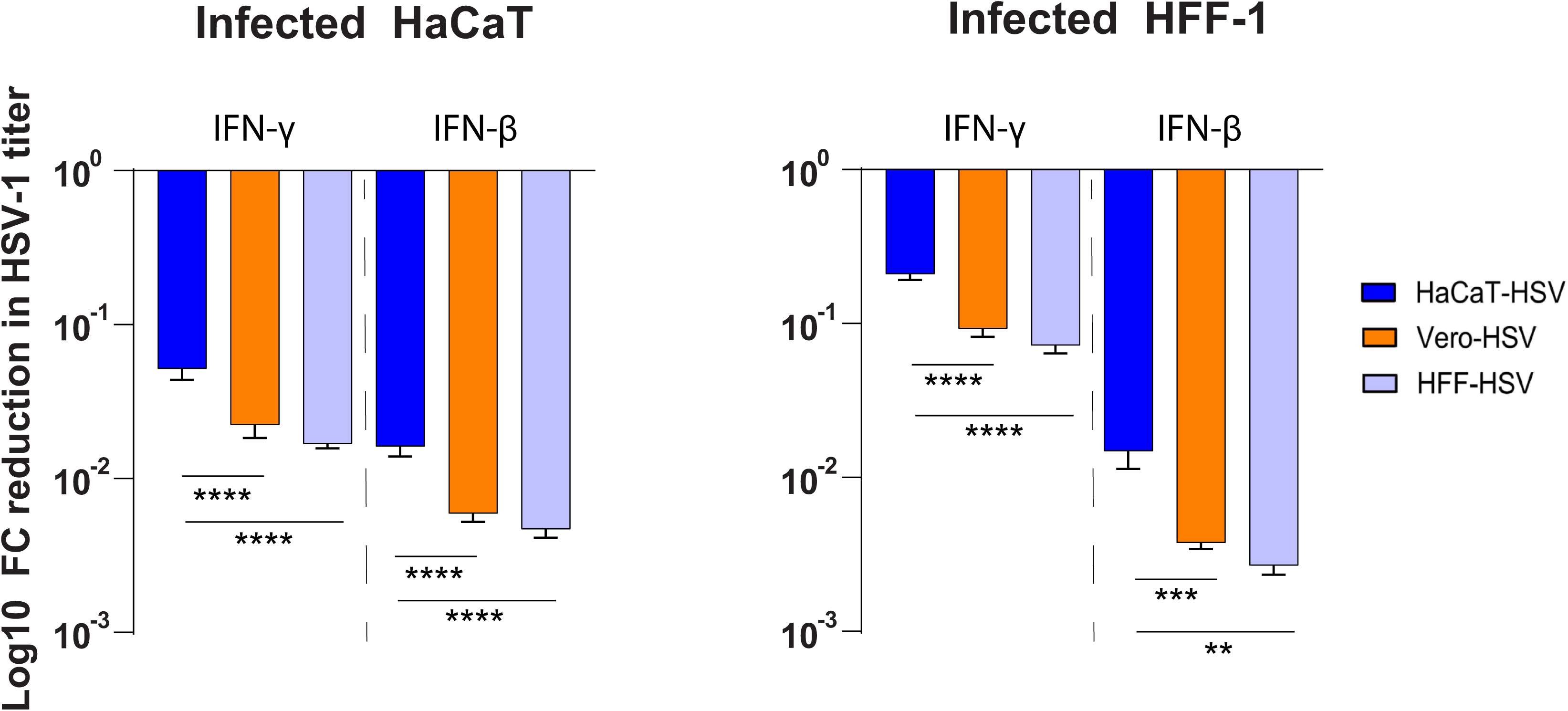
HaCaT-HSV replication is less sensitive to pre-treatment with type I and type II interferon. Cells were pre-treated with 100 U/mL (HaCaT) or 1000 U/mL (HFF-1) for 16 hours prior to infection with HSV-1 McKrae at an MOI of 10. Virally infected cells and supernatant were harvested 12 hpi and titered on vero cells. The values represent the mean reduction in viral titers across two experiments from three biological replicates relative to infected cells that did not receive IFN pre-treatment. Error bars indicate the SEM. Statistical comparison of the producer cell-dependent PFU/cell output between conditions used one-way ANOVA with Tukey’s multiple comparisons test. * indicates p < 0.05, ** indicates p < 0.01, *** indicates p < 0.001, **** indicates p<0.0001.

### Producer cell-dependent effects on the HSV-1 virion proteome

Our results have shown that HSV-1 replication changes based on the producer cell type of HSV-1 virions. HSV-1 expresses 84 viral proteins during infection that perform a wide array of functions critical to viral infectivity and replication^61^. At least 44 of these proteins have been previously identified to be incorporated into mature HSV-1 virions^62–64^. We hypothesized that the phenotypes we have observed are due to producer cell-dependent changes to the proteome associated with HSV-1 virions. To test this hypothesis, HSV-1 McKrae was propagated from HaCaT, vero, and HFF-1 cells and extracellular virus was subjected to a modified version of a previously described purification protocol^64^. Briefly, HSV-1 virions were separated away from cellular components using a combination of centrifugation, filtration, and DNase digestion prior to further purification of virions by ultracentrifugation on a 10% Ficoll cushion. Purified virions were then processed for liquid chromatography and tandem mass spectrometry (LC-MS/MS) of tryptic peptides using a Data Independent Analysis (DIA) workflow. Proteins associated with purified HSV-1 virions were identified and quantified to determine which viral and host cellular proteins are differentially detected between the different samples (Sup. Table 1a). To determine if proteins were either virion-associated or contaminating proteins that co-purify with virions, mock infected cells were scraped and frozen at -80 °C. After thawing, the mock infected cell supernatants were subjected to the same purification scheme, and proteins that were detected from mock-infected cells were identified as potential contaminants and removed from further analysis (Sup. Table 1b-d). As an additional control, separate replicates of HaCaT-HSV virions were concentrated and digested with Proteinase K prior to Ficoll separation^65^. Proteins that are not contained within lipid bilayers are accessible to Proteinase K digestion, resulting in a reduction in the detection of peptides. We considered many of these proteins, other than those known to be virion envelope proteins, to be either a likely contaminant or partial contaminant, meaning that fractions of the protein were both virion and non-virion associated (Sup. Table 2d).

LC-MS/MS identified several viral and cellular proteins that were differentially incorporated into HSV-1 virions based on the producer cell type. 56 total viral proteins were detected in HSV-1 McKrae. Three proteins previously identified as tegument proteins exhibited reduced detection following Proteinase K digestion; thymidine kinase, pUL7, and pUL51 (Sup. Table 2d)^64^. A fourth protein, pUL50 deoxyuridine triphosphatase, was also reduced by Proteinase K digestion, but has not been previously characterized as a tegument protein. Reduced detection suggests that a subset of these proteins are external to the virion envelope. All other Proteinase K accessible viral proteins were envelope proteins, with patterns of reduced detection in line with prior reports of viral membrane proteins^65^. A quantitative comparison of viral proteins showed that 25 and 34 proteins were differentially incorporated into HFF-HSV and Vero-HSV, respectively, relative to HaCaT-HSV (Fig. 4, Sup. Table 2a-c). When applying a stricter threshold for defining differential protein incorporation (greater than 1 log_2_ fold-change with a p<0.01) 39 out of 55 proteins were found to be equivalently detected between HaCaT-HSV and HFF-HSV compared to 26 out of 56 proteins when comparing HaCaT-HSV to Vero-HSV (Fig. 5A). A total of seven viral proteins were detected in HaCaT-HSV and HFF-HSV that were not detected in Vero-HSV. Some of this lack of detection is likely due to the limits of mass spectrometry, including the limited detection of low molecular weight proteins^66^. For example, neither VP26 nor pUs9 were detected in Vero-HSV, and each of these proteins exhibited relatively low or variable detectability between replicate samples in both the HaCaT-HSV and HFF-HSV samples. ICP8 was uniquely detected in Vero-HSV. To address the hypothesis that the virion proteome is contributing to the enhanced infectivity of HaCaT-HSV, we compiled a list of viral proteins that were incorporated into HaCaT-HSV at greater than 1 log_2_ fold-change with a p<0.01 in comparison to both Vero-HSV and HFF-HSV. These proteins included gG, gK, ICP0, and ICP10. We were also interested in viral proteins that may inhibit viral replication. Viral proteins that were incorporated into Vero-HSV and HFF-HSV at greater than 1 log_2_ fold-change with a p<0.01 in comparison to HaCaT-HSV include pUL24 and pUL42.

**Figure 4.**
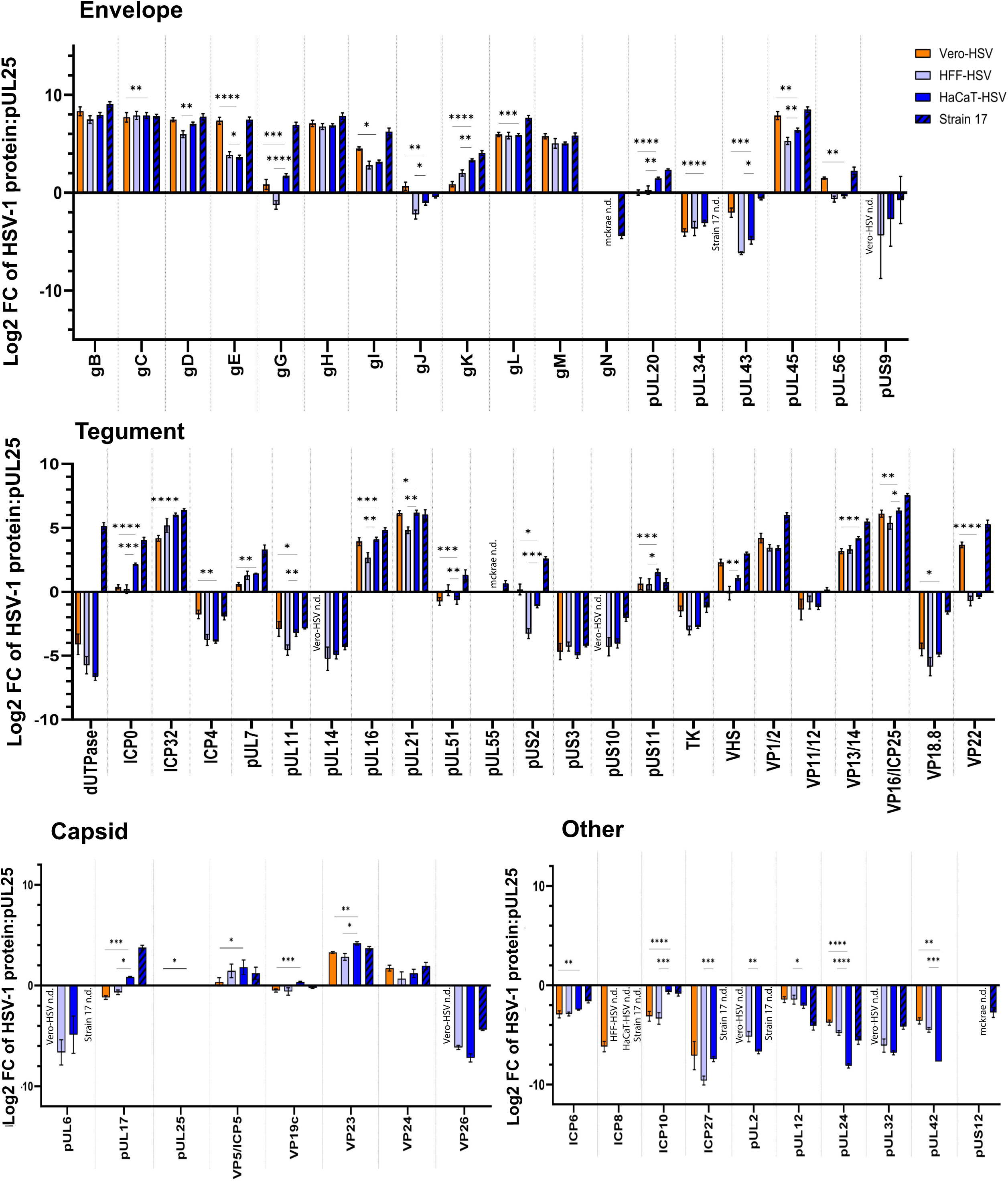
HSV-1 virions incorporate viral proteins in a producer cell and strain-dependent manner. HSV-1 virions were purified from vero and HFF-1 cells (McKrae) or HaCaT cells (McKrae and strain 17). LC-MS/MS was performed on purified virions using an Orbitrap 480 mass spectrometer and protein identification and quantification was based on data-independent acquisition. Relative protein abundance of each viral protein is expressed as the average Log2 fold change (FC) relative to pUL25 detected in each sample type and organized based on the protein localization within the virion. Error bars indicate the SEM within triplicate samples. n.d.: no detection. Statistical comparison of values employed Empirical Bayes statistics for differential detection test. * Indicates p < 0.05, ** indicates p < 0.01, *** indicates p < 0.001, **** indicates p<0.0001

**Figure 5.**
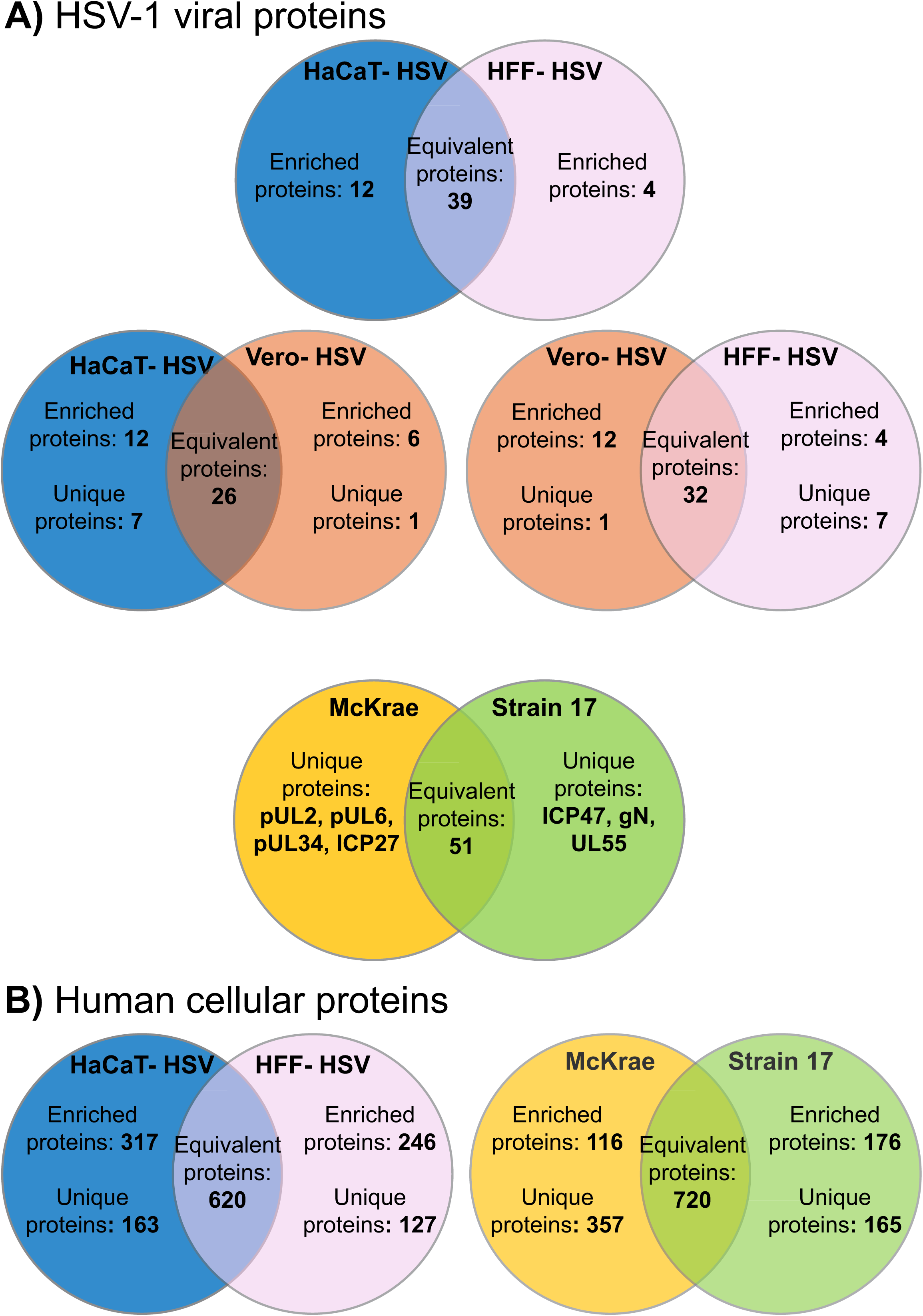
Summary of HSV-1 protein incorporation into virions. Venn diagrams depicit commonalities and differences in HSV-1 viral protein (A) and cellular protein (B) incorporation based on the producer cell type and strain. Strain and cell type being compared are indicated at the top of each diagram.

A quantitative analysis of human cellular protein incorporation into virions was performed comparing incorporation into HaCaT-HSV relative to HFF-HSV (Sup. Table 2b). Comparison of cellular protein incorporation between HaCaT-HSV and Vero-HSV was determined to be unreliable due to protein heterogeneity between species (Sup. Table 2c). Across all samples and controls, 3,225 proteins were identified. A total of 1061 human proteins were detected in both mock infected and HSV-1 infected samples. 185 of these 1061 proteins were detected in greater quantities in viral samples compared to mock. Although some of the cellular proteins identified in mock samples were enriched in viral samples, we decided to treat these as contaminating proteins in the context of our data analysis. Removal of control detected proteins and low-quality detections left 1,880 cellular proteins uniquely detected in HSV-1 infected samples (Sup. Table 3A-C). Additionally, 344 human cellular proteins were accessible to proteinase k digestion in HaCaT-HSV, indicative of either their membrane topology or presence as an extra-virion contaminant (Sup. Table 2D). There were 620 cell proteins that were equivalently detected between HaCaT-HSV and HFF-HSV (Sup. Table 3H). HaCaT derived HSV-1 contained 163 unique proteins, in comparison to 127 proteins unique to HFF-HSV (Sup. Table 3D & 3F). An additional 365 proteins were enriched in HaCaT-HSV relative to HFF-HSV, compared to 283 proteins enriched in HFF-HSV relative to HaCaT-HSV (Sup. Table 3E & 3G). It is difficult to determine what the role of these proteins are in the context of their association with the virion. However, many of the virion-associated cellular proteins are affiliated with vesicular transport, exocytosis, and metabolic processes among others. A summary of the differential detection of viral and cellular proteins based on the producer cell type of HSV-1 can be found in Fig. 5a and 5b.

### Strain dependent effects on the HSV-1 virion proteome

Several previous studies have used LC-MS/MS to identify HSV-1 virion incorporated proteins. These studies have used different viral strains and producer cell types for virion production and analysis (Table 2)^62–64^. Because the incorporation of viral and cellular proteins into HSV-1 virions often varies between studies, we purified HSV-1 strain 17 virions from HaCaT cells to compare to HaCaT-derived McKrae virions.

**Table 2.**
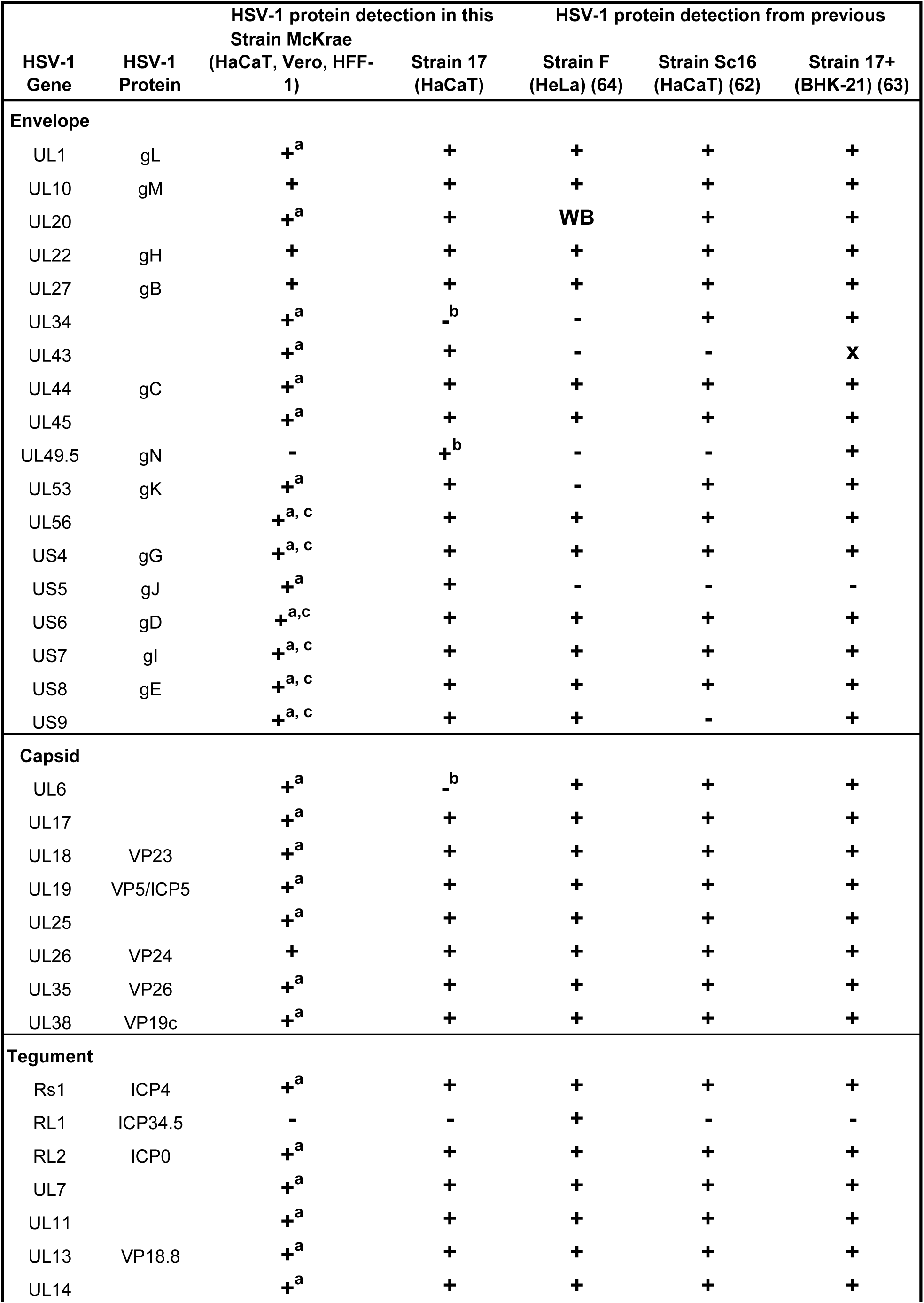
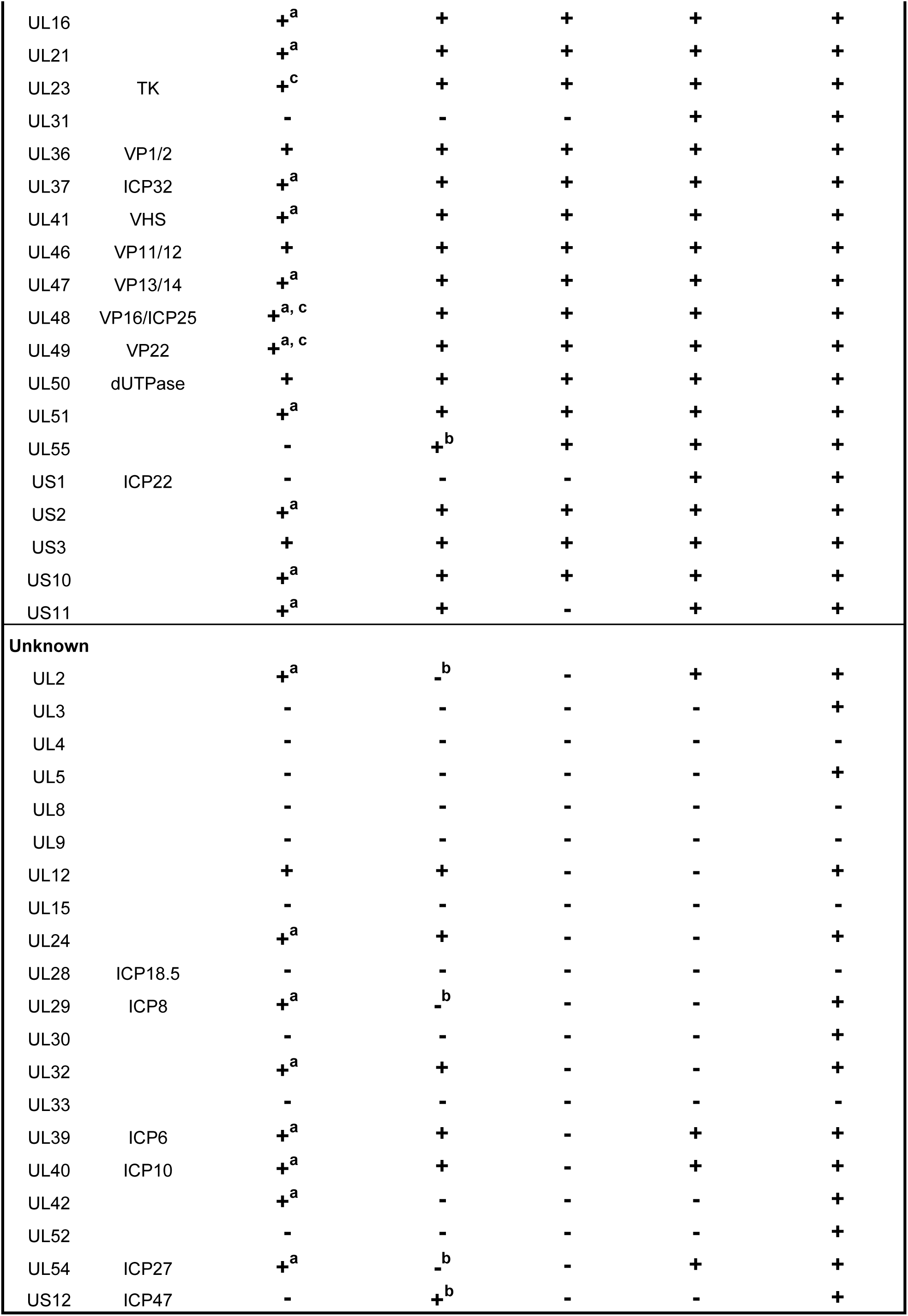
Detection of HSV-1 proteins in purified HSV-1 across studies. Comparison of HSV-1 protein detection by mass spectrometry between our study and previous studies. “+” Detected in purified HSV-1 virions, “-“ not detected in purified HSV-1 virions, superscript “a” producer cell-dependent incorporation in HSV-1 McKrae (current study), superscript “b” strain-dependent effect on viral protein incorporated (current study), superscript “c” the viral protein was accessible to proteinase k digestion (current study), “WB” the viral protein was detected by Western blot but not by mass spectrometry, “x” the viral gene was deleted in the HSV-1 strain used in the study.

A quantitative comparison of HSV-1 McKrae and strain 17 viral protein incorporation was complicated by protein sequence heterogeneity between the two strains. However, LC-MS/MS analysis did reveal a subset of viral proteins that are uniquely incorporated into virions in a strain-dependent manner (Sup. Table 2e). In total, 55 and 54 viral proteins were detected in HSV-1 McKrae and strain 17 derived from HaCaT cells, respectively (Fig. 4). ICP47, gN, and pUL55 were each detected in strain 17 but were undetected in McKrae. Conversely, pUL2, pUL6, pUL34 and ICP27 were all detected in McKrae but not strain 17. Human cellular proteins were abundant in both strains. Strain 17 virions contained 1166 total cellular proteins whereas McKrae incorporated 1361 cellular proteins (Sup. Table 2e). 720 cellular proteins were equivalently detected between each strain, with 181 and 119 of those proteins enriched (greater than 1 log_2_ fold-change with a p<0.05) in strain 17 and McKrae, respectively. Even though each strain was derived from HaCaT cells, 165 unique proteins were detected in strain 17 compared to 357 unique proteins detected in McKrae. The results from our LC-MS/MS analysis show that both the producer cell-type and strain of HSV-1 influences the incorporation of both viral and cellular proteins into virions. A summary of the differential detection of viral and cellular proteins based on the strain can be found in Fig. 5a and 5b.

## Discussion

The producer cell type can alter the infectivity, antigenicity, and cellular tropism of several viruses. Some of these phenotypes have been attributed to producer cell type-dependent changes to the macromolecular contents of the virions themselves. These changes in virion content include altered protein glycosylation patterns, differential protein incorporation, distinct lipid compositions, and selection for genomic mutations. The goal of the work presented here was to determine whether cell type alters the capacity of the produced HSV-1 virions to initiate infection and replicate. The cell types that were selected for these experiments were relevant to HSV-1, including HaCaT and HFF-1 cells. In addition, vero cells were chosen due to their common use as a producer and experimental cell line for HSV-1. The replication capacity of virions from each producer cell type was assessed on multiple cell lines, as measured by viral protein production and the output of infectious virions. When comparing HaCaT-HSV, Vero-HSV, and HFF-HSV infectivity of cells, HSV-1 produced in HaCaT cells exhibited the most robust replication in target cells overall, including cells that had been pre-treated with type I and type II interferon. We hypothesized that the observed differences in HSV-1 replication in target cells are due to producer cell-dependent alterations to the proteome of HSV-1 virions. LC-MS/MS analysis revealed that virions from different producer cells contain unique and differing quantities of both viral and cellular proteins. In addition, HSV-1 McKrae and HSV-1 strain 17 derived from HaCaTs produce unique virion associated proteomes relative to one another. From the data provided by the presented experiments, we can conclude that the producer cell has a direct impact on both the complement of proteins associated with HSV-1 virions and the capacity of virions to replicate. Connecting the changes in the proteome to the producer cell-dependent effects on viral replication will require greater analysis of the individual factors that differ between the samples tested.

Critical to evaluating the effects of a producer cell on HSV-1 is the evaluation of viral protein incorporation. Several viral proteins were differentially detected in both a producer cell-type and strain-dependent manner. 58 viral proteins were identified in purified virions of HSV-1, although one of these proteins, ICP8, was unique to Vero-HSV samples. Viral proteins that were detected at greater quantities (greater than 1 log_2_ fold-change with a p<0.01) in HaCaT-HSV included gG, gK, ICP10, and ICP0. The increased incorporation of ICP0 could explain some of the observed differences in viral replication. ICP0 has multiple activities that include activation of viral gene expression, protein degradation, inhibition of antiviral responses, cell cycle arrest, and inhibition of the DNA damage response^67–73^. Knockout or mutations to ICP0 result in significantly less viral replication in cells that have been pre-treated with type I or type II interferon^73^. In our experiments, HSV-1 derived from each cell type was sensitive to type I and type II interferon in both HaCaT and HFF-1 cells. However, the greater abundance of ICP0 in HaCaT-HSV may have contributed to the reduced sensitivity of HaCaT-HSV infections to interferon pre-treatment. Interestingly, deletion of ICP0 from HSV-1 results in either severely dampened or lack of replication in the cornea and trigeminal ganglia of mice. However, this replication is either partially or fully restored in IFN-α/βR^-/-^ or IFN-α/β/γR^-/-^ mice^72^. Although differential incorporation of viral proteins into virions is not similar to the complete knockout of a viral gene, it does demonstrate the importance of these proteins *in vivo*, and it is possible that producer cell-dependent changes to protein incorporation impacts viral replication and spread *in vivo*. Further testing of the changes to viral protein incorporation will be required to understand if the altered capacity of HSV-1 infection that we have observed is related to the differential incorporation of any of these identified viral proteins.

An alternative interpretation of the cell type differences is that replication is hindered by inclusion of certain viral proteins by Vero-HSV and HFF-HSV. Viral proteins that were incorporated into Vero-HSV and HFF-HSV at greater than 1 log_2_ fold-change with a p<0.01 in comparison to HaCaT-HSV include pUL24 and pUL42. pUL24 and pUL42 each serve many pro-viral roles within infected cells including nuclear remodeling, DNA replication, and the inhibition of antiviral responses^74–79^. Interestingly, both proteins can bind the NF-κB p50 and p65 subunits and prevent the nuclear translocation of NF-κB^80–83^. Although several antiviral roles have been attributed to NF-κB activation, HSV-1 subverts NF-κB function and instead uses it as a pro-viral factor in several cell types including keratinocytes^84–90^. Regions of the HSV-1 genes that encode ICP0 and VP16 each contain κB binding sites, and NF-κB activation is needed for optimal viral gene expression early in infection^87,89,91^. It is possible that the reduced capacity for productive replication by Vero-HSV and HFF-HSV relative to HaCaT-HSV is due to enhanced cytoplasmic sequestration of NF-κB by the pUL24 and pUL42 proteins that are incorporated into virions at greater abundance.

Evaluation of our LC-MS/MS results needs consideration of prior studies that have evaluated HSV-1 virion associated proteomes and the factors that complicate direct comparison. The differential incorporation of proteins by HSV-1 virions based on both the strain and producer cell type may explain why the proteins identified in purified HSV-1 virions often vary between studies^62–64^ (Table 2). As previously mentioned, Yang & Courtney found that HSV-1 strain KOS purified from vero and HEp-2 cells contained ICP0 and ICP4, but not virions derived from BHK21 cells^11^. However, subsequent studies that purified HSV-1 strain 17 or HSV-1 strain F from BHK21 cells were able to detect ICP0 and ICP4 associated with virions^63,64,92^. These conflicting results allude to the possibility that producer cell-dependent effects on HSV-1 virions can change in a strain-dependent manner. Our analysis detected ICP0 and ICP4 in all treatments, although both proteins were incorporated into virions at different quantities in a producer cell-dependent manner.

The detection of virion-associated proteins across studies is further complicated by the use of different methodologies for virion purification and mass spectrometry analysis. These methodological differences may result in variability of protein detection. In addition, mass spectrometers have become far more sensitive and therefore are more likely to detect peptides that may have been undetected in previous studies^93^. Furthermore, it is possible that several of the proteins that have been identified within our LC-MS/MS analysis, as well as other analyses, are not actually associated with HSV-1 virions. Although our purification scheme utilizes several steps to isolate virions away from extra-virion material, our proteinase k-treated sample revealed that several proteins identified by LC-MS/MS were susceptible to digestion. While a subset of these proteins is likely associated with the virion membrane, it is unlikely that this is the case for all proteins that were digested. In addition, we detected several markers that are known to be associated with extracellular microvesicles such as exosomes.

Interpretation of the producer cell-dependent phenotypes must address changes in the complement of cellular proteins that are detected. Due to the large quantity of cellular proteins that were detected in purified virions, it is unclear as to whether these proteins are specifically incorporated into virions or other structures. Stegen et. al previously used siRNA knockdown experiments to determine that a subset of cellular proteins that are incorporated into HSV-1 are important for replication^94^. However, several of these proteins were either not detected in our LC-MS/MS analysis or were incorporated at similar quantities between cell types. Several classes of proteins that were identified have been detected in several viruses, such as translation initiation factors, RNA binding proteins, and cytoskeletal proteins^9,25,62–64,94–103^. Proteins that are known to be associated with exosomes such as tetraspanins, heat shock proteins, and ESCRT proteins are commonly identified in purified virions by LC-MS/MS analysis of several viruses including HSV-1^9,25,62–64,97,100,102,104^. It is possible that extracellular vesicles commonly co-purify with HSV-1 virions, perhaps through aggregation, making it difficult to determine which proteins are truly virion-associated. Previous studies that explicitly isolate exosomes using density ultracentrifugation have not detected HSV-1 DNA within these bands^105,106^. Despite the potential for contamination of our purified virions with extracellular vesicles, it is interesting that the contents of these vesicles may differ based on the producer cell type that is infected. If this is the case, extracellular vesicles excreted during HSV-1 infection may be contributing to the observed differences in HSV-1 replication in infected target cells. Alternatively, it is possible that these proteins that are commonly identified in both exosome and virion purifications are incorporated into HSV-1 virions. A previous study that relied on electron microscopy identified several exosome associated proteins within mature HSV-1 virions^107^. These proteins were specific to exosomes secreted by neurons, none of which have been identified in any previous LC-MS/MS analysis of HSV-1 virions that are isolated from non-neuronal cells. An additional study using a human oligodendroglia cell line identified a fraction of infectious HSV-1 that was secreted in microvesicles^108^. Other studies have determined that HSV-1 directly interacts with exocytosis machinery in non-neuronal cells, providing further evidence that HSV-1 utilizes the exocytosis pathway for egress and as a result, may be incorporating exosome proteins into virions^109^. Indeed, further experiments are needed to validate incorporation of these proteins into virions or to determine whether virions are associated with extracellular vesicles.

An alternative, or perhaps complementary hypothesis as to why the producer cell type of HSV-1 alters the replication capacity of virions is that non-infectious particles produced from each cell line are altering replication. These non-infectious particles can include light particles (L-particles) and defective virions. L-particles produced by HSV-1 infected cells contain viral tegument proteins and an envelope but lack a viral capsid and genome^110^. Previous work has determined that L-particles can function to antagonize immune responses and enhance viral replication of naked HSV-1 DNA^92,111,112^. Inquiries into the nature of L-particles have found that the producer cell type can alter the proteomic contents of L-particles, as well as the ratio of L-particles to mature virions that are produced^11,112^. The role of defective virions in HSV-1 infection requires further research, and a wide array of phenotypes have been observed when studying non-infectious virions in the context of infection for several viruses^29–32^. These roles can include activating or antagonizing antiviral immunity, enhancement or inhibition of viral replication, and induction of cell death depending on the virus or even viral strain that was observed. Our analysis demonstrated that the HSV-1 genome:PFU ratio was lowest in HaCat-derived viral inoculum and that HFF-derived inoculum had the highest genome:PFU ratio. However, inhibitory effects by non-infectious virions could contribute to the differences in HSV-1 replication when the inoculum is derived from different producer cells. Further testing is needed to determine the role of non-infectious virions in the context of HSV-1 infection.

A critical question for these and other studies of HSV-1 virions is the relevance to disease. Studies that have looked at virion associated proteomes of HSV-1 have relied on many cell types, including vero, BHK-21, and HeLa among others, to propagate material for experiments. The use of different producer cell lines for the propagation of viruses can influence experimental outcomes *in vitro* and *in vivo*. For example, the propagation of SARS-CoV-2 and Ross River virus from vero cells results in different outcomes of infection in animal models compared to virus that was propagated from more physiologically relevant cell lines^2,23^. These differential outcomes of infection include altered immune responses, viral dissemination, and a reduction in replication. Future experiments are needed to determine whether the producer cell type of HSV-1 influences infection in animal models. Additionally, HSV-1 can infect many cell types in humans. The spread of HSV-1 primarily begins with replication in keratinocytes associated with the skin or oral mucosa prior to establishing latency in neurons of the peripheral nervous system. In healthy individuals, HSV-1 spread is generally restricted and is either asymptomatic or causes the manifestation of orofacial lesions. However, HSV-1 exhibits a broad tropism in humans resulting in viral replication within tissues of the eye, liver, and central nervous system among others, and replication can even result in systemic spread in some patients^113–116^. Understanding producer cell-dependent effects on HSV-1 may help us to understand viral spread within and between individuals as well as disease outcomes in patients with severe disease.

## Methods

### Cell culture and viruses

HaCaT, vero, and HFF-1 cells were grown in Dulbecco’s modified Eagle’s medium (DMEM) supplemented with 10% fetal bovine serum (FBS) and 1% penicillin/streptomycin (pen/strep) (get actual units) at 37 °C in 5% CO_2_. HSV-1 McKrae was propagated on HaCaT, vero, and HFF-1 cells in DMEM supplemented with 2% FBS and 1% pen/strep. All HSV-1 stocks were tittered by plaque assay on vero cells.

### Fluorescence microscopy

Epifluorescence imaging was performed on a Nikon Ti-Eclipse inverted microscope (Nikon Instruments) equipped with a Spectra X LED excitation module (Lumencor) and fast-switching emission filter wheels (Prior Scientific). Fluorescence imaging used paired excitation/emission filters and dichroic mirrors for DAPI, green fluorescent protein (GFP) and tetramethylrhodamine (TRITC) filters (Chroma corp.). All brightfield images were acquired using phase-contrast configuration at 4x or 20x optical magnification. Imaging immunofluorescent stained samples were set to the same exposure and power when comparing treatments within the same sampling group as determined by cell type and staining time post infection.

### Quantification of HSV-1 antigen

Cells were infected with HSV-1 McKrae using an MOI of 1 or 10. Cells were washed with DPBS 1hpi and replaced with DMEM supplemented with 2% FBS and 1% pen/strep. Cells were fixed at 3 or 6 hpi as described above. Immunofluorescent stains of HSV-1 antigen in infected cells were quantified for fluorescent intensity using Fiji ImageJ. One image from each biological replicate was selected for analysis at each time point. All of the cells within the image were selected as an independent region of interest (ROI). The mean gray value was measured for each ROI and background noise subtracted based on the mean gray value of mock infected cells.

### Purification of HSV-1 virions

HSV-1 McKrae was purified from HaCaT, vero, and HFF-1 cells using a modified purification scheme that was previously described (Lippe). Cells were infected with HSV-1 McKrae at an MOI of 0.01. Inoculum was removed 1 hpi. Cells were washed with DPBS and replaced with serum-free DMEM. HSV-1 was propagated for 48 hrs (HaCaT) or 72 hrs (vero and HFF-1). Viral supernatants were harvested and centrifuged at 300 xg for 10 min. The supernatant was then filtered through a 0.45 μm filter. Virions were pelleted by centrifugation at 20,000 xg for 30 min at 4 °C. Virions were resuspended in PBS, sonicated, and incubated with 100 U/mL of DNAse I at 37 °C for 1 hr. Virions that received proteinase k treatment were incubated with 0.1 mg/mL of Proteinase K for 45 min at room temperature and subsequently treated with 2 mM of phenylmethylsulfonyl fluoride (PMSF). Samples were further diluted in DPBS, loaded onto a 10% ficoll cushion, and centrifuged at 26000 x RPM for 2 hrs at 4 °C on an SW41 Ti rotor. The pellets were resuspended in 100 uL of 0.75% NP-40 for 1 hr on ice. Purified HSV-1 was stored at -80 °C. This same protocol was used for HSV-1 strain 17 purified from HaCaT cells and mock-infected HaCaT, vero, and HFF-1 cells that were scraped and frozen at - 80 °C prior to purification.

### HSV-1 genome quantification using qPCR

HSV-1 genome was concentrated and isolated to form a standard curve for qPCR quantification of HSV-1 viral genomes from viral stocks. To do this, HSV-1 supernatants were harvested from HaCaT cells infected with HSV-1 McKrae at an MOI of 10 for 20 hrs. Supernatants were centrifuged at 300 xg for 10 min and filtered through a 0.45 μm filter. Virions were pelleted by centrifugation at 20,000 xg for 30 min at 4 °C. Virions were resuspended in PBS, sonicated, and incubated with 100 U/mL of DNAse I at 37 °C for 1 hr to remove contaminating cellular DNA. HSV-1 genome was then isolated from digested virions using the Zymo Quick-DNA miniprep kit (cat# D3024). The HSV-1 viral genome concentration was calculated using nanodrop prior to building the standard curve for qPCR analysis.

HSV-1 virion-associated genome quantitation was calculated by comparing HSV-1 genome content to known standard curve values of HSV-1 genomic DNA. Prior to qPCR analysis, HSV-1 viral stock was treated with DNAse I and processed through the Zymo Quick-DNA miniprep kit as previously described. qPCR analysis was performed using PowerUp SYBR Green kit (Applied Biosystems, cat no. 4367659), targeting HSV-1 ICP27. 28S rRNA was targeted as quality control to ensure no cellular DNA was present in standard curve or sample DNA.

qPCR primers:

28S fwd 5’- GGG CCG AAA CGA TCT CAA CC −3’

28S rev 5’- GCC GGG CTT CTT ACC CAT T −3’

ICP27 fwd 5’- AGA CGC CTC GTC CGA CGG −3’

ICP27 rev 5’- AGG CGC GAC CAC ACA CTG T −3’

### Fixation and immunofluorescent staining

Cells were washed with DPBS and fixed with ice-cold 50:50 methanol:acetone for 20 minutes at RT. Methanol:acetone was removed, dried, and cells were blocked with 2% bovine serum albumin (BSA) in DPBS for 10 min at RT. Cells were incubated with rabbit polyclonal HSV-1 anti-serum (1:500, Dako) diluted in antibody solution (1% BSA in DPBS) for 1 hr at RT. Cells were washed 3 x with DPBS and incubated with Donkey anti-Rabbit IgG Alexa Fluor 488 (2 drops/mL) diluted in antibody solution for 1 hr at RT. Cells were washed with DPBS and incubated with 1:5000 hoescht in DPBS for 10 min. Cells were washed 2 x with DPBS prior to fluorescent imaging.

### Plaque assay

Cells were infected with HSV-1 McKrae at an MOI of 10. The inoculum was removed 1 hpi and cells were washed with DPBS and incubated in cell media consisting of DMEM supplemented with 2% FBS and 1% pen/strep at 37 °C in 5% CO_2_. Cells and supernatants were harvested by scraping at corresponding time points. Samples were stored at -80 °C and sonicated after thawing for measurement of viral titers by plaque assay on vero cells. For experiments in which interferon pre-treatment was used, cells were treated with 100 U/mL (HaCaT) or 1000 U/mL (HFF-1) of interferon beta, gamma, or both. Interferon was removed from cells after 16 hours of incubation, washed with DPBS, and infected as described above.

### Liquid chromatography-mass spectrometry

Total protein from each sample was reduced, alkylated, and digested using single-pot, solid-phase-enhanced sample preparation with sequencing grade modified porcine trypsin (Promega)^117^. Tryptic peptides were trapped and eluted on 3.5um CSH C18 resin (Waters) (4mm x 75um) then separated by reverse phase XSelect CSH C18 2.5 um resin (Waters) on an in-line 150 x 0.075 mm column using an UltiMate 3000 RSLCnano system (Thermo). Peptides were eluted at a flow rate of 0.300uL/min using a 60 min gradient from 98% Buffer A (0.1% formic acid, 0.5 % acetonitrile):2% Buffer B (0.1% formic acid, 99.9% acetonitrile) to 95:5 at 2.0 minutes to 80:20 at 39.0 minutes to 60:40 at 48.0 minutes to 10:90 at 49.0 minutes and hold until 53.0 minutes and then equilibrated back to 98:2 at 53.1 minutes until 60 minutes. Eluted peptides were ionized by electrospray (2.4 kV) through a heated capillary (275°C) followed by data collection on an Orbitrap Exploris 480 mass spectrometer (Thermo Scientific).

Precursor spectra were acquired with a scan from 385-1015 Th at a resolution set to 60,000 with 100% AGC, max time of 50 msec, and an RF parameter at 40%. DIA was configured on the Orbitrap 480 to acquire 50 x 12 Th isolation windows at 15,000 resolution, normalized AGC target 500%, maximum injection time 40 ms)^118^. A second DIA was acquired in a staggered window (12 Th) pattern with optimized window placements. Following data acquisition, data were searched using Spectronaut (Biognosys version 19.1) against the UniProt *Homo sapiens*, UniProt *Chlorocebus sabaeus,* UniProt HHV1 strain17, and NCBI HHV1 McKrae (April 2024) respectively, using the directDIA method with an identification precursor and protein q-value cutoff of 1%, generate decoys set to true, the protein inference workflow set to maxLFQ, inference algorithm set to IDPicker, quantity level set to MS2, cross-run normalization set to false, and the protein grouping quantification set to median peptide and precursor quantity. Protein MS2 intensity values were assessed for quality using ProteiNorm^119^. The data was normalized using VSN and analyzed using proteoDA to perform statistical analysis using Linear Models for Microarray Data (limma) with empirical Bayes (eBayes) smoothing to the standard errors^120–125^. Proteins with an FDR adjusted p-value < 0.05 and a fold change > 2 were considered significant.

## Acknowledgements and Funding

The mass spectrometry analysis was performed in collaboration with the IDeA National Resource for Quantitative Proteomics at the University of Arkansas, funded by NIH/NIGMS grant R24GM137786. We are thankful for the work performed by all members of the Resource, in particular Dr. Samuel Mackintosh and Dr. Dennis Province for their patience with our many questions during consultation and analysis.

